# Few Effects of a 5-week Computerized Cognitive Training Program in Healthy Older Adults

**DOI:** 10.1101/570143

**Authors:** Sheida Rabipour, Cassandra Morrison, Jessica Crompton, Marcelo Petrucelli, Murillo de Oliveira Gonçalves Germano, Anita Popescu, Patrick S. R. Davidson

## Abstract

Computerized cognitive training programs are becoming increasingly popular and practical for cognitive aging. Nevertheless, basic questions remain about the benefits of such programs, and about the degree to which participant expectations might influence training and transfer. Here we examined a commercial cognitive training program (*Activate*) in a 5-week double-blind, pseudo-randomized placebo-controlled trial. Based on a priori power analysis, we recruited 99 healthy older adults 59-91 years of age (M = 68.87, SD = 6.31; 69 women), assigning them to either the intervention or an active control program (Sudoku and n-back working memory exercises). We subdivided both groups into high and low expectation priming conditions, to probe for effects of participants’ expectations on training and transfer. We assessed transfer using a battery of standard neuropsychological and psychosocial measures that had been agreed to by the training program developers. We planned and pre-registered our analyses (on osf.io). The majority (88%) of participants progressed through the training, and most provided positive feedback about it. Similarly, the majority (80%) of participants believed they were truly training their brains. Yet, transfer of training was minimal. Also minimal were any effects of expectations on training and transfer, although participants who received high expectation priming tended to engage more with their assigned program overall. Our findings suggest limited benefits of *Activate* training on cognition and psychosocial wellbeing in healthy older adults, at least under the conditions we used.

## Introduction

Subtle decline in several cognitive domains is typical even in healthy aging (Anderson & Craik, 2017; Davidson & Winocur, 2017; Drag & Bieliauskas, 2010; Toepper, 2017). To prevent or slow this cognitive decline, computerized interventions are increasingly becoming popular among researchers, clinicians, and older adults themselves. Such Computerized Cognitive Training (CCT) represents an attractive alternative to medication and other interventions that may be costly and have serious side effects (National Academies of Sciences, 2017). It is therefore unsurprising that older adults are prominent targets of – and subscribers to – commercialized CCT programs (e.g., Lumosity, Cognifit, Brain Age; Shah, Weinborn, Verdile, Sohrabi, & Martins, 2017).

Nevertheless, basic questions remain about CCT. An obvious one is whether such programs actually work. When people ask this question, they are usually asking about transfer of training: That is, will training lead to improvement on tasks that were not trained, but that rely on the same underlying cognitive processes as those that underwent training? Answering this question has been difficult due to the poor methodological quality of previous research on CCT. For example, studies have often suffered from insufficient control conditions – notably, small sample sizes, frequent lack of an active control group, and inadequate consideration of potential placebo effects (W. R. Boot, Simons, Stothart, & Stutts, 2013; Green et al., 2019; Motter, Devanand, Doraiswamy, & Sneed, 2016; National Academies of Sciences, 2017; Simons et al., 2016). Importantly, cognitive training trials have frequently employed passive (e.g., waitlist) controls as a comparison for the training group(s). This practice is problematic as it permits only an evaluation of test-retest effects without providing any insight into nonspecific factors related to participating in a program.

Here we report a trial of the commercial CCT program *Activate* in older adults, in which we addressed many of these limitations, including the largely ignored potential confound of participant expectations. In particular, we aimed to evaluate the real-world practicality of such training by examining transfer effects, beyond expected practice (i.e., test-retest) effects (Collie, Maruff, Darby, & McStephen, 2003), using standard neuropsychological measures.

### Activate Program

Although many commercial CCT programs are available (for a partial list and review, see Simons et al., 2016), we selected the *Activate* web-based program because of its focus on training working memory (WM), sustained and divided attention, inhibition, and visuospatial perception. Many of these functions decline even in healthy aging (Davidson & Winocur, 2017; Reuter-Lorenz, Festini, & Jantz, 2016; Zanto, Chadick, & Gazzaley, 2014). Specifically, the *Activate* games are divided into four categories, targeting: i) spatial WM; ii) visual pursuit and category sorting; iii) visual pursuit and inhibition; and iv) “pattern completion” (i.e., determining the image that best fits into an empty space, to complete a pattern).

In our view, the *Activate* program had two advantages: First, the training is adaptive. In other words, the program adjusts for individual performance on each task, remaining at a certain level of difficulty as long as participant accuracy and response times continue to improve at a certain rate. The program only increases level of difficulty once performance has reached a plateau, rather than advancing to the next level after a predetermined number of correct trials. This approach allows participants to master each task at a given level of difficulty before moving on to subsequent levels. Researchers have argued that such adaptive training methods are particularly likely to produce transfer of training (Heinzel et al., 2014; Lovden, Backman, Lindenberger, Schaefer, & Schmiedek, 2010; Morrison & Chein, 2011). Second, the *Activate* program had already shown promise of transfer in older adults with depression, with training reported to reduce depressive symptoms and measurably improve cognitive control (as indexed by the Trail-Making and Stroop tests; Morimoto et al., 2016; Morimoto et al., 2014). A recent report further demonstrated the potential of *Activate* to improve children’s performance on executive function measures in the classroom setting (Kavanaugh, Tuncer, & Wexler, 2018). Despite these promising preliminary reports, no other evaluations of this program currently exist (Simons et al., 2016). Thus, we sought to replicate these previous reports by including the neuropsychological tasks from those studies (i.e., Trail-Making and Stroop tests), and came to a consensus with the *Activate* developers on other tasks that should show transfer in older adults.

### Expectations & Placebo Effects

Much of the CCT literature has been criticized for having relatively low methodological quality (e.g., Motter et al., 2016; Simons et al., 2016). Perhaps the most serious problems revolve around insufficient control conditions, including lack of an appropriate active control group to distinguish intervention effects from those of placebo or other non-specific factors, and failure to account for participants’ expectations of training outcomes (W. R. Boot et al., 2013; Foroughi, Monfort, Paczynski, McKnight, & Greenwood, 2016). Previous surveys have suggested that older adults have optimistic expectations of CCT (Rabipour, Andringa, Boot, & Davidson, 2017; Rabipour & Davidson, 2015). Thus, what appear to be transfer effects following CCT could, instead, merely represent placebo effects. To our knowledge, expectations have rarely – if ever – been explicitly addressed in CCT trials (for the few existing examples, see W. R. Boot et al., 2013; Foroughi et al., 2016). A powerful way to accomplish this is with a balanced-placebo design (Rohsenow & Marlatt, 1981), using a randomized double-blind placebo-controlled design and sub-dividing both the treatment and placebo control groups such that half of participants in each group are told they are in the treatment condition and the other half told they are in the placebo control group. Such a design helps one tease apart the true effects of the intervention from those of expectations.

### The Present Study

Here we examined a 5-week CCT program in healthy older adults actively seeking preventative cognitive interventions. Using a balanced-placebo design, we aimed to elucidate the potential influence of such expectations on cognitive performance following participation in CCT. We took particular care in our study design and data collection to ensure: i) reasonable statistical power; ii) use of a dynamic CCT program adaptive to participant performance; iii) use of an active (rather than less-rigorous passive) control condition (Redick, 2015; Shipstead, Redick, & Engle, 2010), comprised of a computerized program that would believably pass as cognitive training; iv) control of potential expectancy-mediated placebo effects (W. R. Boot et al., 2013; Green et al., 2019); v) use of consensus transfer measures that the developers of the CCT program agreed should reflect the cognitive processes being trained by their exercises, and that are standard measures used in both clinical and experimental studies, including in previous evaluations of the *Activate* program; vi) replication and extension of previous findings on *Activate* improving executive function in older adults (Morimoto et al., 2014), and of our own work on older adults having higher expectations of CCT (Rabipour & Davidson, 2015); and vii) double-blind design. These steps follow best practice recommendations (e.g., Green et al., 2019; National Academies of Sciences, 2017).

## Methods

### Participants

We recruited 99 healthy older adults (M age=68.88, SD=6.31 years; 69 women) with normal or corrected-to-normal vision and hearing from the Ottawa community via ads and flyers (Figure 1). We screened all participants based on the following exclusion criteria: i) diagnosis of memory impairment or cognitive decline; ii) diagnosis of neurological disorder; iii) experience of stroke or transient ischemic attack; and iv) experience of injury or operation resulting in long-term changes in memory or cognitive function.

A priori power analyses using G*Power version 3.1 determined that a minimal sample of 84 participants (i.e., 21 older adults per group) would be required for repeated-measures ANOVA with within-between subjects interactions, based on a moderate effect size (Cohen’s f(V)=.45) at 80% power and alpha=.01 to account for multiple measurements. This interaction between assigned group (see below) x time (pre vs. post training) – and, in particular, between time and training program – would provide the strongest evidence of *Activate* having specific transfer effect(s).

**Figure 1.**
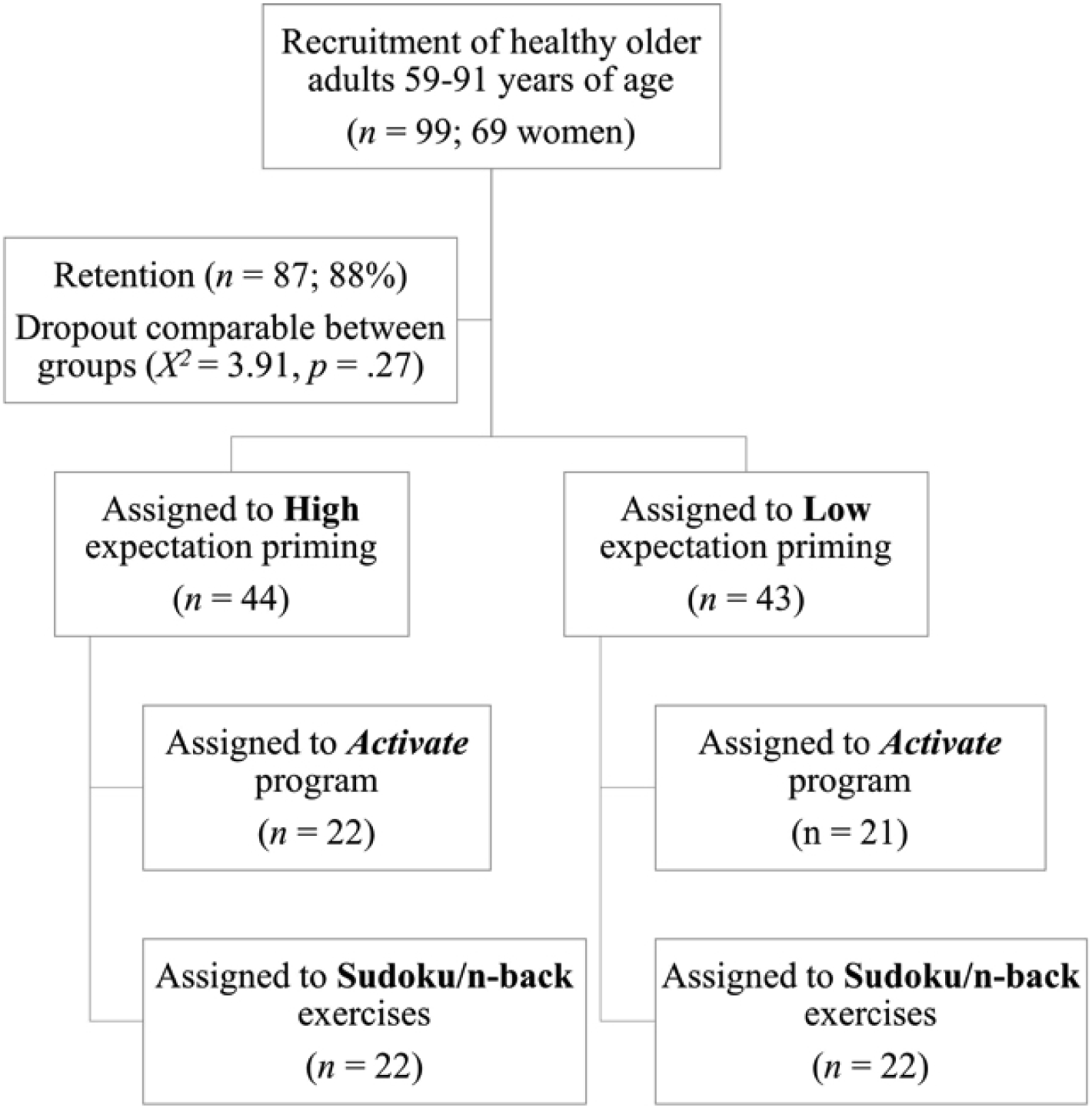
Participant recruitment and group assignment with final sample.

### Experimental Design and Protocol

The study was a double blind randomized controlled trial (registered at clinicaltrials.gov, ID: NCT0220571), crossing expectation priming (high vs. low) with CCT program (*Activate* vs. placebo-controlled Sudoku and n-back exercises). After providing consent, participants were assigned to one of two expectation priming conditions: i) High expectation priming, in which participants were told they would receive a type of CCT known to improve performance; and ii) Low expectation priming, in which participants were told they would receive a type of CCT with no known benefits. We then assigned participants to receive one of two programs: the commercialized *Activate* CCT intervention, or an active control (see below).

We followed a pseudo-randomization procedure, aiming to balance the age and sex of participants across groups as best as possible. Importantly, our expectation priming conditions and comparison of two separate CCT programs enabled blinding of participants with respect to their experimental condition. To avoid divulging the nature of the study and potentially skewing the results, participants who entered the study with a friend or family member, or who were referred to the study by a past participant, were assigned to the same group. To maintain blinding for researchers, we ensured that those examining participants would not be present during the training sessions and would have no contact with participants outside the pre-and post-training assessments.

Both CCT programs comprised five 40-minute training session per week, over five weeks. The 25 sessions each involved two 20-minute periods of training, separated by a brief break (i.e., several minutes) if desired. Each week, participants were instructed to complete a minimum of four home sessions in addition to one lab session, carried out in a small group setting at the University of Ottawa main campus (i.e., where several participants would complete their training together in the same room, on individual computers). The rationale for the weekly lab sessions twofold: First, the supervised lab sessions helped us ensure that participants were adhering to the correct training protocol and were not experiencing any issues related to the program or understanding of the training tasks. Second, previous research has suggested that older adults benefit from CCT to a greater extent when completing the program, at least in part, in supervised group sessions conducted at a center (Lampit, Hallock, & Valenzuela, 2014; Wadley et al., Dividing the home and lab sessions in this manner permitted us to balance between convenience and quality control. Similarly, we selected the 5-week time frame to promote practicality and minimize participant attrition, while permitting a reasonable amount of training (i.e., 1000min or ±17h), comparable to other training studies and plausibly leading to transfer (e.g., Kavanaugh et al., 2018).

We administered behavioural assessments at baseline and following completion of the CCT program. We also performed a partial long-term follow-up assessment, wherein we asked participants to complete select questionnaires six months following the completion of the program. At the time of this long-term follow-up, participants were contacted via telephone to provide final comments on their experience and perspective on the effects of the program via an informal interview.

In order to maximize intrinsic motivation and avoid possible performance decrements associated with receipt of extrinsic rewards (Jaeggi, Buschkuehl, Shah, & Jonides, 2014), participants received no monetary or other compensation for participation in the CCT program. However, participants were able to retain free access to the program beyond their participation in the study. Moreover, to compensate for time spent in the research lab at the University of Ottawa, participants were offered free parking as well as a chance to enter a lottery to win a 50$CAD gift card. We received ethical approval to conduct this study from the Research Ethics Boards at the University of Ottawa and the Bruyère Research Institute.

### Expectation Assessment and Priming

Participants rated their expectations of CCT effectiveness using the Expectation Assessment Scale (EAS), a validated questionnaire (Rabipour, Davidson, & Kristjansson, 2018) we have previously used to measure and prime expectations with written messages implying that the intervention is either highly effective or not expected to produce any meaningful change (see Rabipour, Andringa, et al., 2017; Rabipour & Davidson, 2015). We administered the EAS on three occasions: i) at baseline; ii) after participants received the expectation priming messages (High or Low); and iii) after completing the 5-week CCT program. Participants rated their expectations of outcomes on a scale from 1-7 (1 = “completely unsuccessful,” 2 = “fairly unsuccessful,” 3 = “somewhat unsuccessful,” 4 = neutral/“I have absolutely no expectations,” 5 = “somewhat successful,” 6 = “fairly successful,” 7 = “completely successful”). We probed expectations on seven cognitive domains: i) general cognitive function; ii) memory; iii) concentration; iv) distractibility; v) reasoning ability; vi) multitasking ability; and vii) performance in everyday activities. Following the training program, we probed participants about their experience using a feedback questionnaire (see supplemental files).

### Training Programs Activate training

The intervention, the pirate-themed *Activate* web program (C8 Sciences), comprised six adaptive computer games involving WM, sustained and divided attention, inhibition, and visuospatial perception. Specifically, the games are divided into four categories based on targeted cognitive functions: i) spatial WM (two games); ii) visual pursuit and category sorting (one game); iii) visual pursuit and inhibition (two games); and iv) pattern completion (one game).

The *Activate* intervention aims to account for individual differences in performance on each task: each task remains at a level of difficulty as long as participant accuracy and response times continue to improve, rather than advance after a predetermined number of correct trials. The program increases level of difficulty only once performance reaches a plateau. Originally created to enhance school performance in children with impulse-control impairments, the program has more recently been applied to older adults and appears to reduce depressive symptoms as well as executive dysfunction in patients with geriatric depression (Morimoto et al., 2016; Morimoto et al., 2014).

During each session, participants were presented with a choice of three possible games. The selected game was played for five minutes, after which the program terminated the game and presented participants with a new choice of three games. Once participants completed four 5-minute game blocks (i.e., 20 minutes of training), the program automatically logged participants out. We asked participants to complete two 20-minute training periods in immediate succession (with the possibility of a brief break in between), at least five different days per week, resulting in five weekly 40-minute sessions in total. Although participants were able to select which set of games to play throughout each session, the program limited choice if participants strongly favoured certain games over others, to ensure roughly equivalent amount of training with all games.

### Sudoku and single n-back training

The active control program comprised non-adaptive computer games with no unifying theme. Participants had a choice of two game types: Sudoku or single n-back WM exercises. We selected these games based on popular conceptions that they broadly improve cognitive functions (Walter R. Boot et al., 2016), coupled with little empirical evidence supporting this notion (Shipstead, Redick, & Engle, 2012; Souders et al., 2017). Unlike the intervention groups, participants assigned to the active control condition selected games entirely at their own discretion for each session. We nevertheless encouraged participants to split their sessions evenly between Sudoku puzzles and n-back games. Similar to the *Activate* training, participants were automatically logged out of the game portal after 20 minutes of task engagement, and were asked to complete two 20-minute training blocks in immediate succession (again, with possibility of a brief break), at least five different days per week.

For the Sudoku puzzles, participants had the option to select “easy”, “medium”, “hard”, or “evil” levels, as per traditional Sudoku. While we wanted to ensure adequate engagement with the Sudoku puzzles, we also sought to further reduce the possibility that this control program would enhance cognitive function or benefit participant performance on untrained tasks. Thus, based on pilot testing, the Sudoku puzzles included in all levels beyond “easy” were, in reality, randomized to generate an easy-level puzzle on 80% of trials, and a puzzle of the selected level on the remaining 20% of trials. Moreover, the “evil” level was, in reality, equivalent to “hard” (i.e., we did not actually include any “evil” level Sudoku puzzles in our program). Notably, because the control program was intended solely as a comparison rather than training activity, we prioritized minimizing difficulty while maintaining engagement. Our analyses of these data reflect the level of Sudoku participants actually completed, rather than the level they selected.

For the n-back tasks, participants had the option of selecting 2-, 3-, or 4-back tasks. Meta-analyses suggest minimal evidence of transfer, especially for single-domain n-back (Redick & Lindsey, 2013), whereas our goal was to evaluate far transfer effects. The type of n-back was randomized to comprise: i) a randomly generated string of letters and numbers; ii) a randomly generated string of pictures; or iii) a spatial n-back task involving a cartoon brain image randomly moving across a 3×3 grid. Due to technical issues with our database, we were unable to retain data regarding level of n-back for the majority of our participants; we analyzed these data based on n-back type.

### Transfer Measures

We included screening measures of global cognition and language function at baseline as potential discriminating variables in our subsequent analyses. We assessed global cognition using the Montreal Cognitive Assessment (MoCA), a brief cognitive screening tool sensitive to mild declines in cognitive function (Nasreddine et al., 2004). Based on advice from our colleagues, we also used the 15-item Boston Naming Test (Lansing, Ivnik, Cullum, & Randolph, 1999) as a screen for language function in our final 53 participants.

As indicators of transfer, we assessed performance on commonly used neuropsychological tests and subjective reports of cognition, quality of life (QOL), and affect. Here we focus on the neuropsychological tests (for the other data, see supplemental files). These measures were agreed on *a priori* by the lead authors (S.R. and P.S.R.D) and the creators of the *Activate* program as reasonable indicators of transfer:

#### Wisconsin Card Sorting Task (WCST)

The WCST, which requires participants to categorize cards by colour, shape, or number using only computer feedback, is a measure of executive function and cognitive flexibility (Rhodes, 2004). We evaluated number of categories completed, an index sensitive to age-related changes in executive function (Rhodes, 2004).

#### California Verbal Learning Test, second edition (CVLT-II)

The CVLT assesses verbal memory by presenting the participant with two lists of 16 words, divided into four semantic categories, over several trials. The full version of the CVLT, used in the present study, involves a short-and long-delay component separated by 15-25 minutes. Scores are typically lower with increasing age and lower level of education, as well as in men compared to women (Lamar, Resnick, & Zonderman, 2003). We selected to evaluate the semantic clustering (Stricker, Brown, Wixted, Baldo, & Delis, 2002) and long-delay cued recall sub-components of the CVLT, which index strategy use (i.e., organizational clustering) and recall ability, respectively.

#### Digit Span Task

The digit span task, a subtest of the Wechsler Adult Intelligence Scale, is an index of WM capacity. For the forward digit span (FDS) task, a measure of short-term recall, participants repeat a string of numbers that the examiner reads aloud; in the backward digit span (BDS) task, an index of WM capacity, participants recite the string of numbers in the reverse order. In the present study, participants completed both the FDS and BDS tasks, but only scores for the BDS were included in our analyses as a measure of executive function (Glisky, Polster, & Routhieaux, 1995).

#### Trail Making Test (TMT)

The TMT requires participants to draw a straight line between numbers (Trail A) and alternating numbers and letters (Trail B) in sequential order, as quickly as possible. The time taken to complete each trail provides a measure of visuospatial capacity, executive control, and cognitive flexibility (Kortte, Horner, & Windham, 2002). Here we administered both Trails A and B, but included only Trail B time for our final analyses, as per previous studies of *Activate* (Morimoto et al., 2016; Morimoto et al., 2014).

#### Controlled Oral Word Association Test (COWAT) & Animal Naming Test

The COWAT measures letter fluency (LF) with the generation of words that begin with a given letter within 60 seconds. Participants name as many words as possible that begin with the letter “F”, then “A”, and finally “S”. The animal-naming test, administered immediately after the COWAT, is a measure of semantic fluency that requires participants to name as many animals as possible in 60 seconds. Scores represent the total number of admissible words reported and appear sensitive to age-related changes in verbal fluency. Although participants completed both the LF and animal naming tasks, only scores for the LF (analyzed as an average number of words per letter) were included in our analyses as an index of executive function (Glisky et al., 1995; McCabe, Roediger, McDaniel, Balota, & Hambrick, 2010).

#### Stroop Interference Task

One of the most commonly used measures in cognitive psychology, the Stroop task (Stroop, 1935) measures attention, processing speed, inhibition, and executive control. The test proceeds in three timed parts of 45 seconds each: In the first part, which measures reading speed, the participant is given a list of colour words and must read as many words as possible within the allotted time. In the second part, which measures colour-naming speed, the participant must name the ink color of printed “X”s as quickly as possible. In the final part – the interference condition – the participant must quickly name the color of the ink in which the colour words are written, without reading the words themselves (e.g., say “blue” when seeing the word “green” written in blue ink). The version used in the present study contains 120 words per part; scores in each part of the test are combined to create an interference score. Because it is brief, requires very little education, and is not culturally biased, this test is a useful screen for neuropsychological deficits. Older adults appear to have slower interference scores, which may reflect decreased efficiency of inhibitory processing (Spieler, Balota, & Faust, 1996).

#### Letter-Number Sequencing

In a subset of our participants (n=69), we additionally explored performance on letter-number sequencing, a subtest of the Wechsler Adult Intelligence Scale. This brief, standardized measure of executive function indexes verbal and visuospatial WM performance, as well as processing speed (Crowe, 2000). During the task, examiners read a series of scrambled letters and numbers to participants, who must then recite the numbers in sequential order, followed by the letters in alphabetical order.

### Statistical Analysis

We performed analyses using IBM SPSS Statistics, Inc. version 24, R version 3.1.1, and JASP version 0.9.2. We pre-registered our analysis plan on Open Science Framework, as part of the “OSF Pre-registration Challenge”.

We conducted repeated-measures multivariate ANOVA (MANOVA) on each category of outcomes (i.e., expectation ratings, training tasks, and neuropsychological and psychosocial transfer measures) with expectation priming and CCT conditions as between-subjects factors, and time as a within-subjects factor. We followed up significant results with simple effects ANOVA and t-tests between the groups of interest, and non-significant results with Bayesian repeated measures ANOVA using the default parameters in JASP. Where applicable, we used the Holm-Bonferroni correction for multiple comparisons and Greenhouse-Geisser correction for sphericity.

## Results

MANOVA between assigned program (*Activate* vs. control) and expectation priming condition (high vs. low) indicated no significant group differences in age, years of education, cognitive function (MoCA and Boston Naming scores), or medication use (Wilk’s λ≥.867, F_(5,35)_≤1.08, p≥.39, BF_10_≤.479; Table 1). Chi-squared analysis similarly revealed no significant group differences in sex (*X* =1.06, p=.787). Self-reports of medication use indicated that the majority of participants (78/87=90%) did not take anticholinergic medication. Moreover, the distribution of Anticholinergic Cognitive Burden scores (Campbell, Maidment, Fox, Khan, & Boustani, 2013) did not significantly differ between groups (*X*^2^ =2.31, p=.511).

**Table 1.** Participant demographics. Groups did not significantly differ with respect to distribution of sex, mean age, years of education, global cognition and memory, language function, and self-reported number of medications ingested.

| Group | Program | Expectation Priming | *n*<br>(women) | Age | Years of Education | MoCA Global Score | MoCA Memory Score | Boston Naming Test | Medication Number |
| --- | --- | --- | --- | --- | --- | --- | --- | --- | --- |
| 1 | *Activate* | High | 22 (17) | 67.59<br>(6.89) | 16.55<br>(2.48) | 27.5<br>(2.69) | 3.23<br>(1.82) | 14.55<br>(.52) | 2.11 (1.84) |
| 2 | *Activate* | Low | 22 (14) | 69.59<br>(6.91) | 17.64<br>(3.53) | 26.86<br>(2.77) | 2.45<br>(2.02) | 13.82<br>(.98) | 3.00 (3.25) |
| 3 | Sudoku/n-back | High | 21 (14) | 68.95<br>(4.44) | 15.86<br>(2.97) | 27.29<br>(2.45) | 3.24<br>(1.79) | 14.45<br>(.82) | 2.29 (2.37) |
| 4 | Sudoku/n-back | Low | 22 (15) | 69.64<br>(7.22) | 16.77<br>(2.41) | 26.48<br>(3.19) | 3.00<br>(1.90) | 14.18<br>(.98) | 2.41 (2.50) |

Participant retention rate was high: after dropout due to scheduling conflicts (n=5), health issues (n=4), loss of interest (n=2), and other personal emergencies (n=1), 88% (87/99; 60 women) of our sample completed the study and were included in our final analyses.

### Expectations of Outcomes

Participants in all groups were largely optimistic of CCT outcomes at baseline (Figure 2), with ratings in all cognitive domains significantly above neutral for participants in all groups (t_21_≥2.11, p≤.047, Cohen’s d≥.45), with the exception of ratings for “multitasking” in participants assigned to *Activate* training and low expectation priming (t_21_=.84, p=.41, BF_10_=.031). Mean expectation ratings did not significantly differ across groups at baseline.

**Figure 2.**
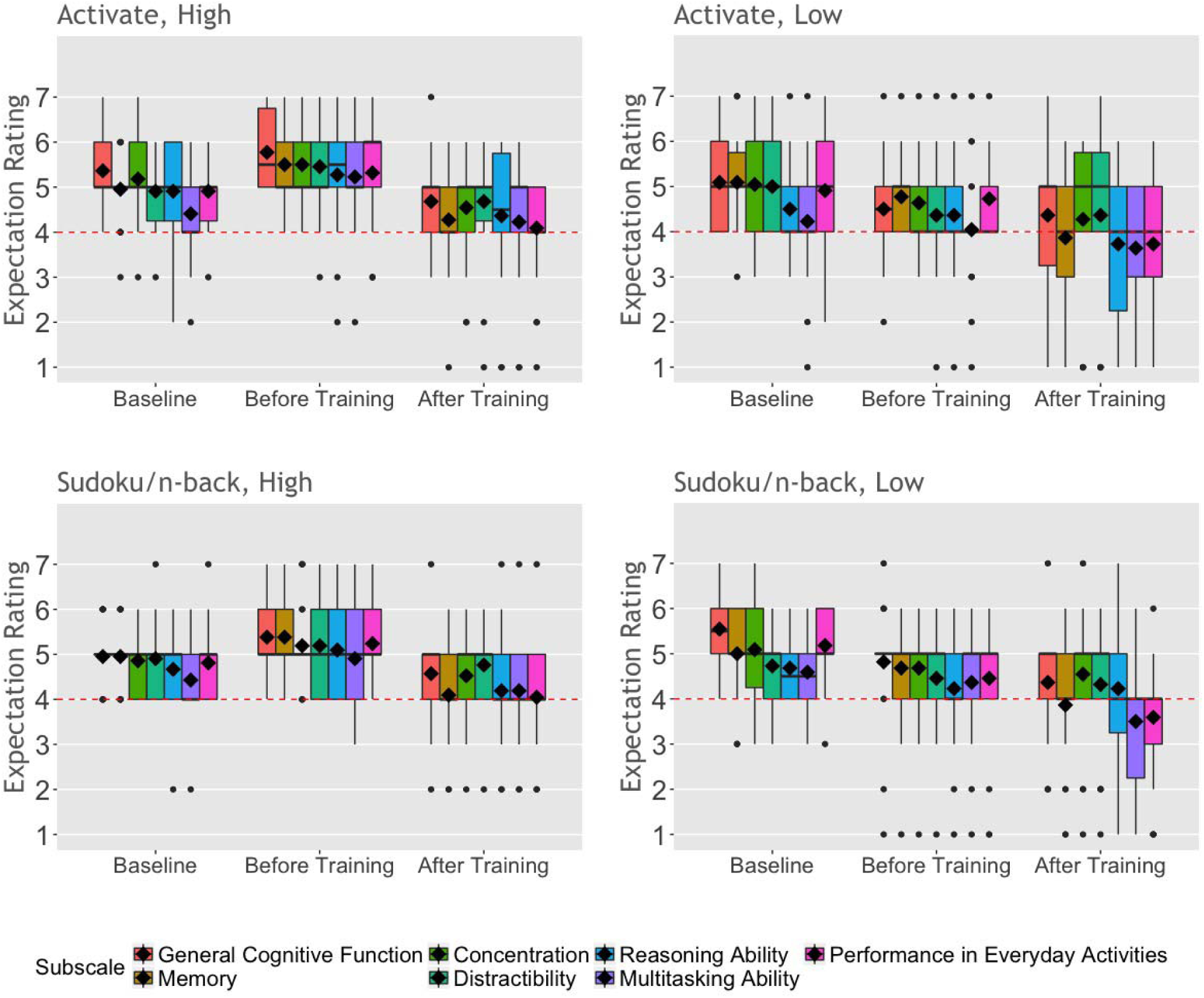
Participant EAS ratings at baseline, after receiving expectation priming (i.e., before training), and following completion of their assigned 5-week CCT program. Ratings of expected outcomes were on a scale of “1” (“completely unsuccessful”) to “7” (“completely successful”) for each subscale (i.e., cognitive domain). Red dashed lines represent a neutral rating of “4” (“no expectation”). Solid lines represent group medians; diamonds represent group means.

We performed repeated measures MANOVA comparing expectation ratings across the seven cognitive domains between expectation priming condition and CCT program at three times: baseline, after receiving the expectation priming message, and after completing the CCT. This analysis revealed a significant main effect of time (Wilk’s λ=.349, F_(14,70)_=9.31, p<.0001, η_p_^2^ =.65) as well as a significant interaction between time and expectation condition (Wilk’s λ=.618, F_(14,70)_=3.08, p=.001, η_p_^2^ =.38; Figure 2). We did not find a main effect of program or of expectation condition, or an interaction effect, on expectation ratings in any cognitive domain, after correcting for multiple comparisons (BF_10_≤.906).

Probing the results for each cognitive domain, we found a significant effect of time (F_(1.58,131.4)_≥11.17, p<.0001, η_p_^2^ ≥.104) and significant interaction between time and expectation condition (F_(1.58,152.15)_≥3.50, p≤.044, η_p_^2^ ≥.04) for all cognitive domains, except for “distractibility”, where the effect of time (F_(1.58,130.7)_=3.30, p=.051, η_p_^2^=.038; BF_10_=.786) was marginal.

Pairwise comparisons showed that participants in the high expectation condition significantly increased their expectation ratings immediately after receiving the priming message, compared to baseline (i.e., from baseline to before training) in all cognitive domains (M_diff_ ≥.33; p≤.003). Conversely, after correcting for multiple comparisons, participants who received low expectation priming decreased their expectation ratings only for the “general cognitive function” domain (M_diff_ ≥.66; p≤.0001), after receiving the low expectation priming message. After correcting for multiple comparisons, we did not find group differences in ratings for any cognitive domain following completion of the program.

### Training Performance

#### Engagement with Training Tasks

Using repeated measures MANOVA, we evaluated the amount of time participants in each expectation priming condition engaged in the *Activate* games over the five weeks (Figure 3a). We found a main effect of week (Wilk’s λ=.309, F_(16,27)_=3.77, p=.001, η_p_^2^=.69) and of expectation priming (Wilk’s λ=.762, F_(4,39)_=3.04, p=.028, η_p_^2^=.24), but no interaction between expectation and week (F_(16,27)_=1.63, p=.128, BF_10_=.051). Further examination revealed the effect of week to be significant for the “visual pursuit / category sorting” (F_(4,168)_=2.93, p=.022, η_p_^2^=.065) and “visual pursuit / inhibition” (F_(4,168)_=13.81, p<.0001, η_p_^2^=.25) categories, and the effect of expectation to be significant for the “visual pursuit / category sorting” games (F_(1,42)_=5.76, p=.021, η_p_^2^=.12). Specifically, participants who received high expectation priming engaged significantly more with the “visual pursuit / category sorting” games, compared to those in the low expectation priming condition. Regarding the effect of week, participants engaged in the “visual pursuit / category sorting” games significantly more during the fourth week of the program, compared to the first, second, and fifth weeks (t_43_ ≥2.27, p≤0.028, Cohen’s d≥.69); for the “visual pursuit / inhibition” category, participants engaged in the games significantly less in the fourth and fifth weeks (t_43_ ≥2.67, p≤0.011, Cohen’s d≥.81), compared to the first three. Based on these results, we included amount of time played per week as a covariate in analyses of performance outcomes (see below).

**Figure 3.**
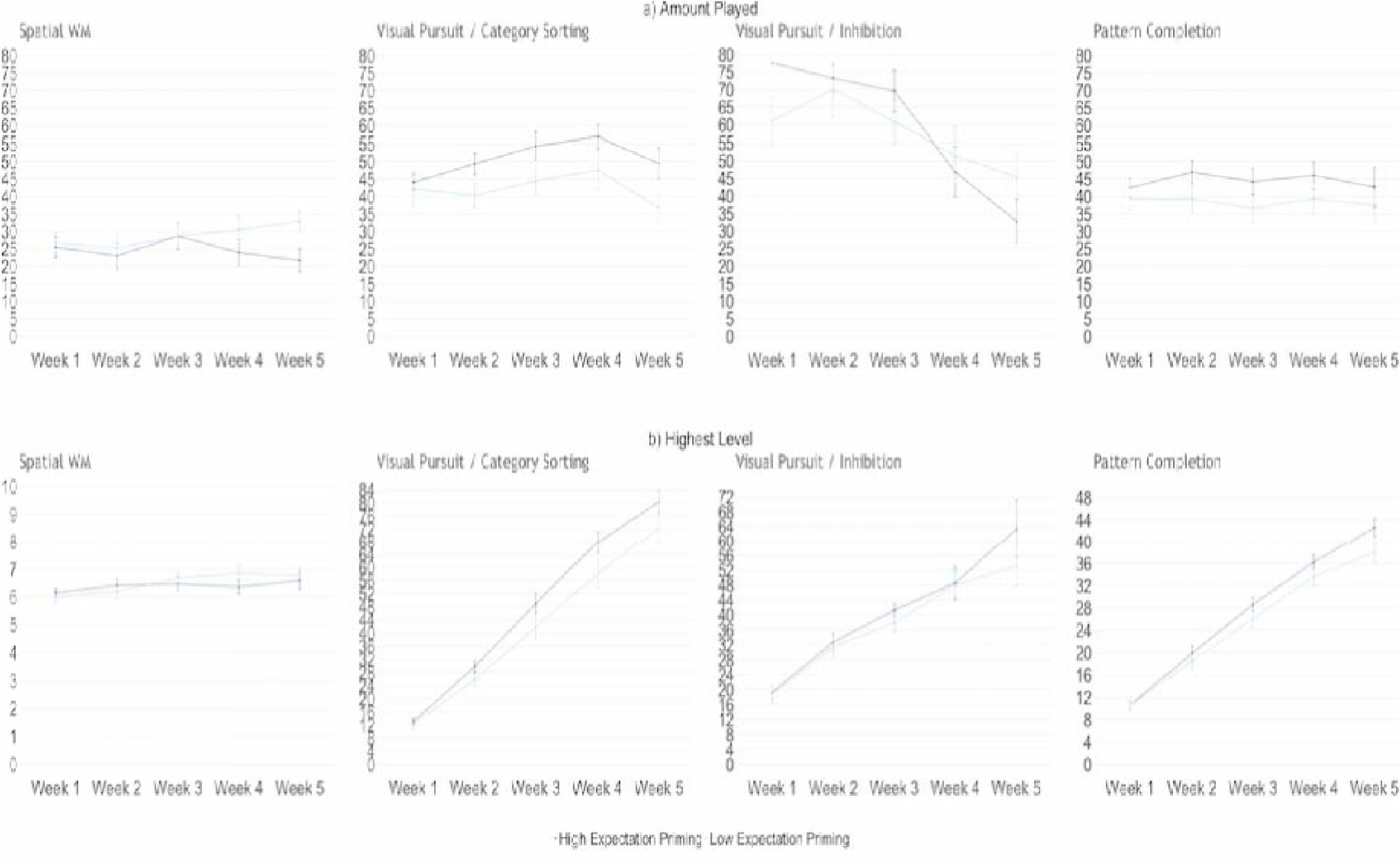
Engagement and performance on the *Activate* games represented by a) the amount of time (min) spent on each game and b) the highest level attained on each game. Error bars represent SEM.

Similarly, for participants in the control program, we used repeated measures MANOVA to examine amount of engagement in each level of Sudoku puzzle and type of n-back exercise over the five weeks (see supplemental files). We found no significant effect of time (Wilk’s λ=.22, F_(24,12)_=1.77, p=.152, BF_10_=.012), of expectation priming (Wilk’s λ=.955, F_(6,30)_=.24, p=.961, BF_10_=.117), or interaction between time and expectation priming (Wilk’s λ=.307, F_(24,12)_=1.13, p=.43, BF_10_=.003).

### Performance on Training Tasks

Because we found differences in amount of engagement with the program, we included total amount of time played (i.e., sum of played amount over the five weeks) as a covariate in our repeated measures multivariate analysis of performance on the *Activate* training tasks (Figure 3b). We found a main effect of time on highest level achieved in the *Activate* games (Wilk’s λ=.371, F_(16,25)_=2.64, p=.014, η_p_^2^ =.63), but no main effect of expectation priming (F_(4,37)_=1.90, p=.132, BF_10_=.301) or interaction between time and expectation priming (F_(16,25)_=.91, p=.56, BF_10_=.01). Follow-up univariate tests revealed the effect of time to be significant for the “spatial WM” (F_(2.7,107.1)_=4.25, p=.009, η_p_^2^ =.096) and “visual pursuit / category sorting” categories (F_(1.5,59.9)_=6.65, p=.005, η_p_^2^ =.14). For the “spatial WM” games, where levels represent list length of items remembered, performance was significantly lower in the second week, compared to the first (p=.041), fourth (p=.029), and fifth weeks (p=.006); for the “visual pursuit / category sorting” games, performance steadily and significantly improved over the five weeks (p<.0001). Note that visual inspection suggests improved performance on the “visual pursuit / category sorting”, “visual pursuit / inhibition”, and “pattern completion” games over the five weeks. However, based on results form our MANCOVA, these patterns are likely influenced by the amount of time spent on the games.

We performed the same analysis to evaluate performance on the control tasks in two separate MANOVAs to account for the two different n-back scores (hit and false alarm rates) analyzed (see supplemental files). For both the number of successfully completed Sudoku puzzles and performance on n-back games, we found no effect of time (Wilk’s λ=.828, F_(12,30)_=.519, p=.89, BF_10_=.023; Wilk’s λ=.373, F_(24,18)_=1.26, p=.31, BF_10_=6.837×10^-4^, respectively), of expectation priming (Wilk’s λ=.938, F_(3,39)_=.866, p=.47, BF_10_=2.729; Wilk’s λ=.977, F_(6,36)_=.144, p=.99, BF_10_=.309, respectively), or an interaction between the two (Wilk’s λ=.652, F_(12,30)_=1.33, p=.25, BF_10_=.002; Wilk’s λ=.493, F_(24,18)_=.773, p=.726, BF_10_=.002, respectively).

### Neuropsychological Transfer Effects

Repeated measures MANOVA examining performance on neuropsychological transfer tests across expectation condition and program revealed a main effect of time (Wilk’s λ=.633, F_(7,75)_=6.22, p<.0001, η_p_^2^=.37) and of expectation priming (Wilk’s λ=.761, F_(7,75)_=3.36, p=.004, η_p_^2^=.24; Figure 4). Notably, we found no significant main effect of program (BF_10_≤.961) or evidence of interactions between time, program, and expectation priming (BF_10_≤1.892).

**Figure 4.**
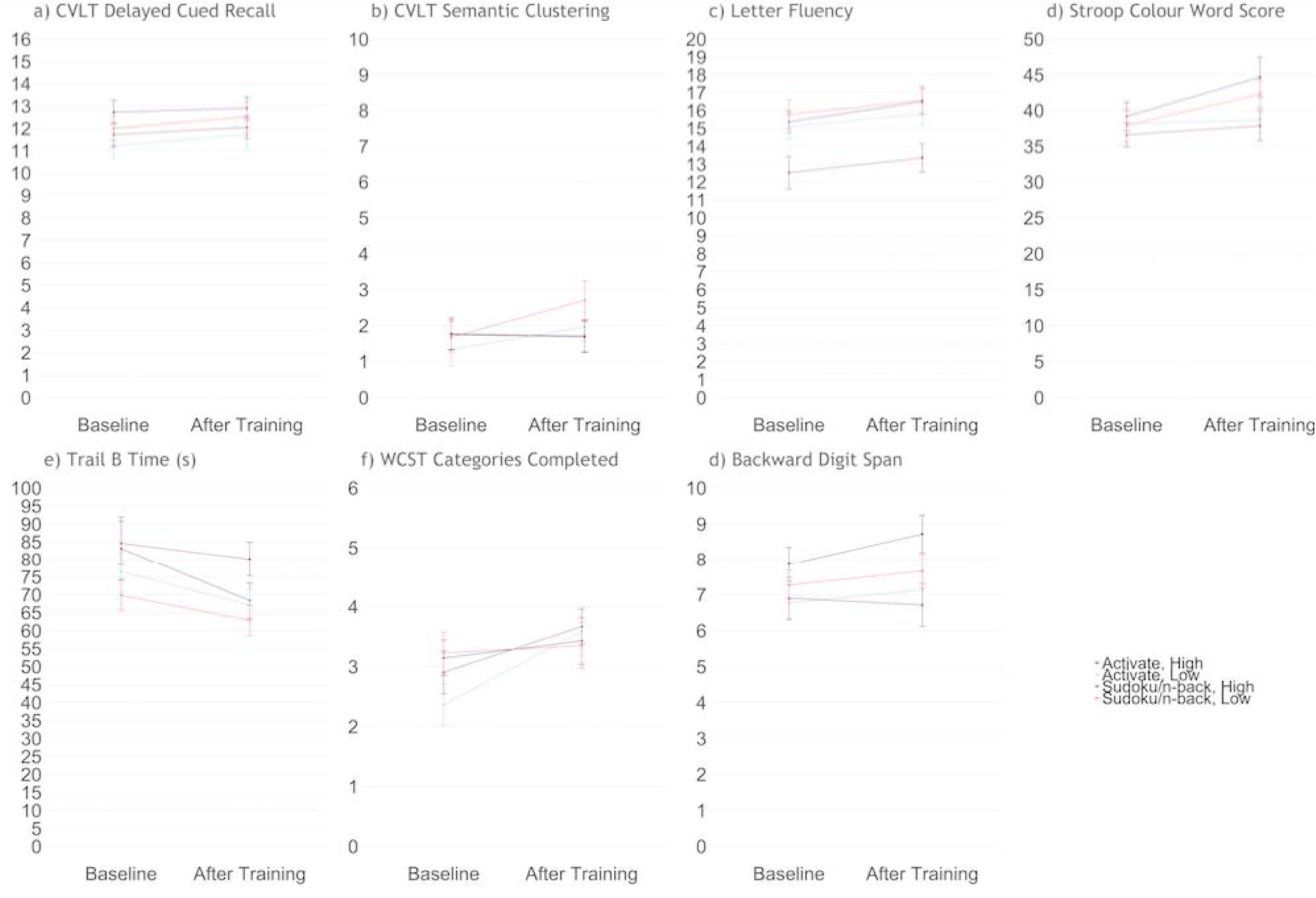
Performance on cognitive tasks analyzed, including: a) long-delay cued recall and b) semantic clustering on the CVLT; c) letter fluency, computed as the average number of words beginning with the letters F, A, and S; d) Stroop colour-word score; e) time (s) on Trail B of the TMT; f) categories completed on the WCST; and g) score on the BDS. Error bars represent SEM.

**Figure 6.**
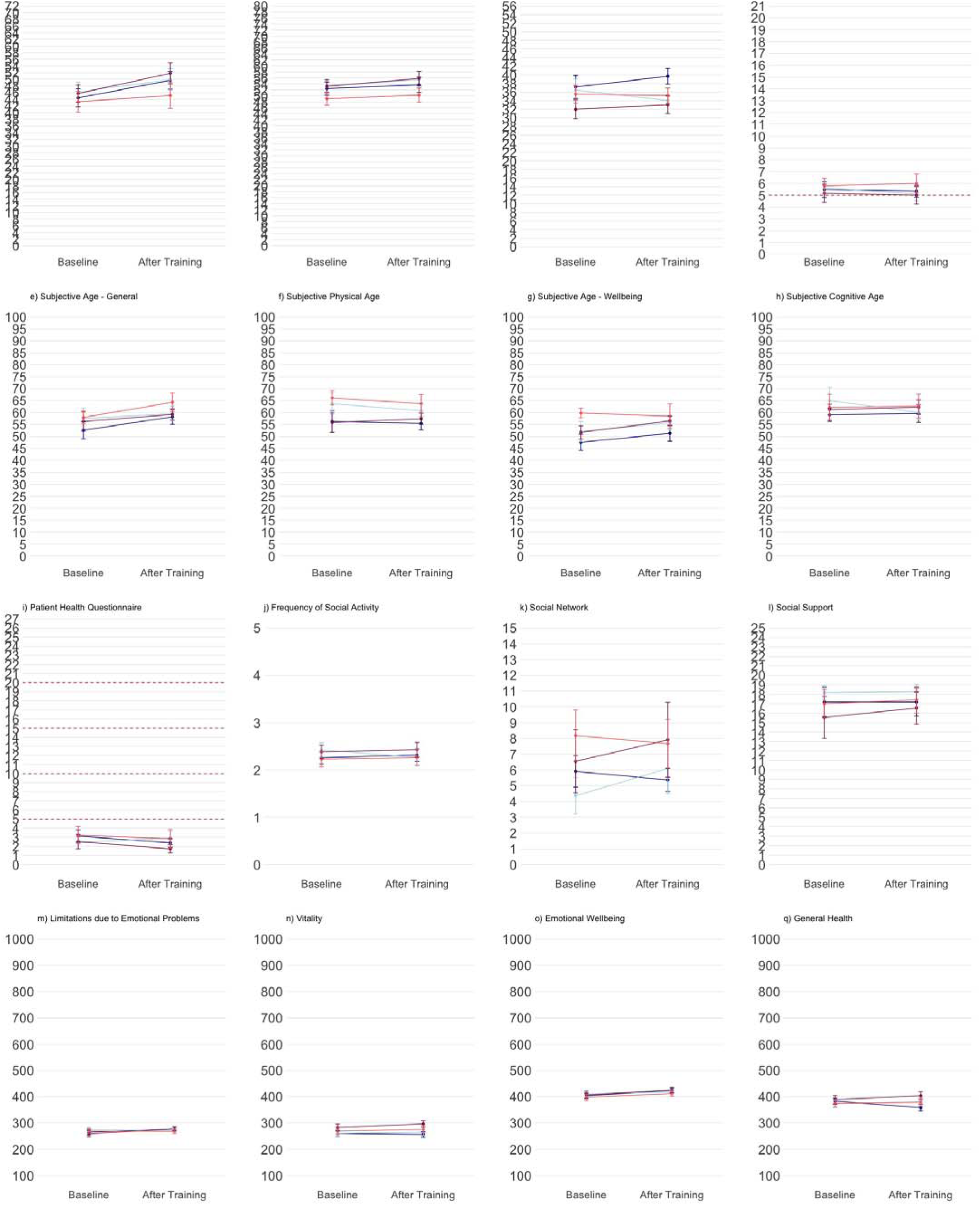
Subjective reports of (a-c) memory, (d) sleep quality, (e-h) perceived age, (i) mood, (j-l) social engagement, and (m-p) quality of life. Solid blue lines represent Activate groups; solid red lines represent control groups. Darker lines represent high expectation priming groups; lighter lines represent low expectation priming groups. Dashed red lines represent clinical threshold(s), where applicable. Error bars represent SEM.

**Figure 7.**
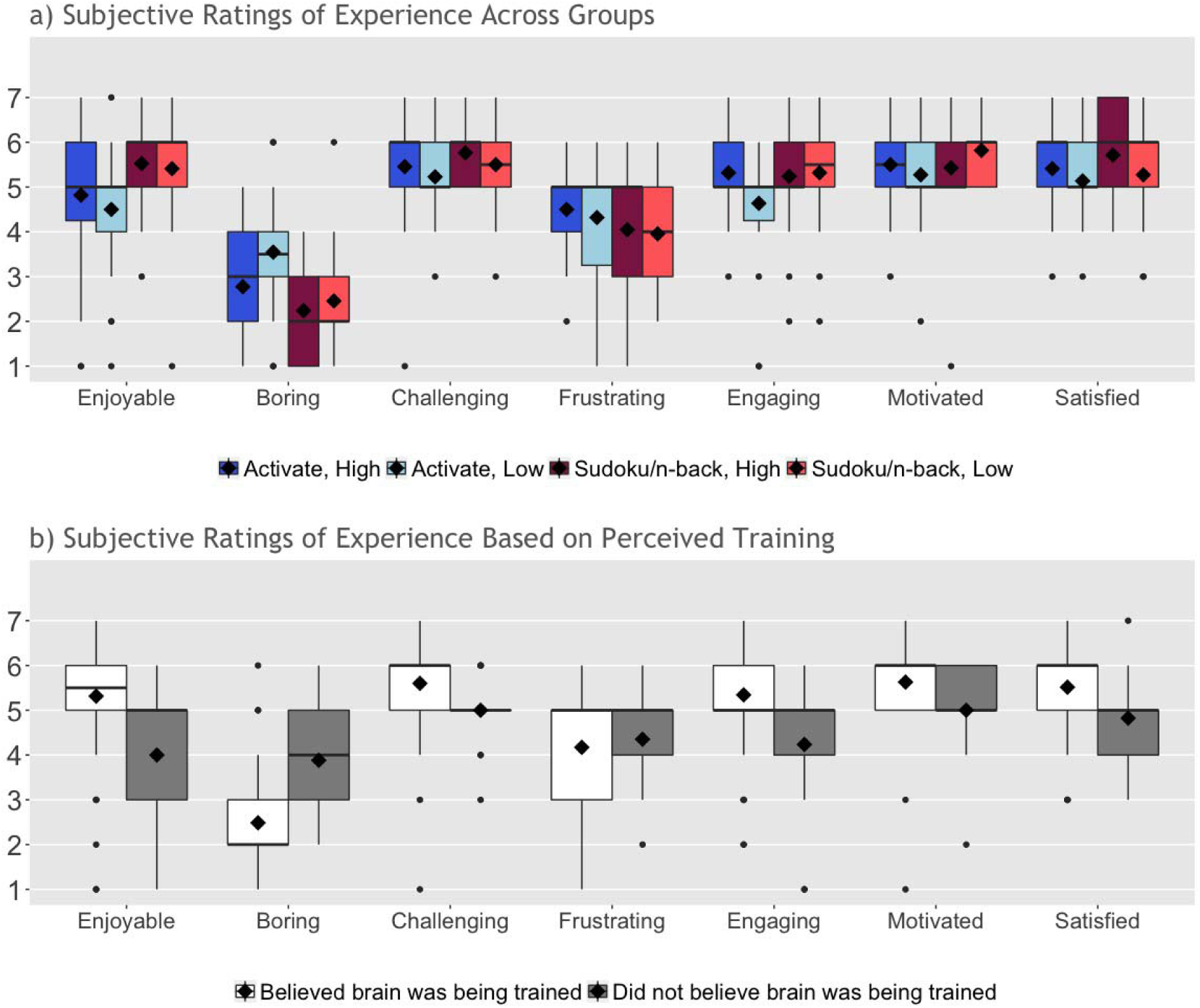
Subjective feedback on experience based on a) experimental group and b) belief that the assigned program genuinely trained the brain. Feedback ratings were on a scale of “1” (“completely disagree”) to “7” (“completely agree”) for each subscale. Solid lines represent group medians; diamonds represent group means.

Follow-up analyses indicated a significant post-training improvement in Stroop colour-word score (F_(1,81)_=11.35, p=.001, η_p_^2^=.12), time on TMT trail B (F_(1,81)_=10.87, p=.001, η_p_^2^=.12), categories completed on the WCST (F_(1,81)_=9.88, p=.002, η_p_^2^=.11), letter fluency (F_(1,81)_=7.20, p=.009, η_p_^2^=.08), and CVLT semantic clustering (F_(1,81)_=4.24, p=.043, η_p_^2^=.05). Moreover, the effect of expectation was significant for time on TMT Trail B (F_(1,81)_=4.61, p=.035, η_p_^2^=.05), whereby participants assigned to low expectation priming had faster response times.

## Discussion

Here we sought to evaluate the effects of expectation priming and a 5-week CCT program on cognitive function and psychosocial wellbeing in healthy older adults. Most participants adhered to their assigned program (88%), believed they were genuinely training their brains (80%) and reported a positive experience overall. All participants improved somewhat on the neuropsychological and psychosocial transfer measures over time, but training on the *Activate* program yielded no greater improvement on the transfer measures than the control training. Similarly, the high expectation priming condition led to no greater improvement (on training or transfer) than the low expectation condition.

### Participant Expectations

As in our previous work (Rabipour, Andringa, et al., 2017; Rabipour & Davidson, 2015), here we found that older adults were largely optimistic about CCT outcomes before they began the study. Moreover, ratings at the outset of the program suggested that our expectation priming (i.e., via the EAS) had successfully raised or lowered participant expectations. Following completion of the program, however, participants were more uncertain about the effectiveness of the CCT in improving cognitive outcomes, with ratings closer to the “neutral” score in all experimental groups.

### Activate and Control Training

Participants assigned to the *Activate* CCT adhered to the program and improved in three out of four of the games’ training domains, including “visual pursuit & category sorting”, “visual pursuit & inhibition”, and “pattern completion”. The relative lack of improvement in the “spatial WM” exercises may reflect the restricted range of these game levels, which represented list length remembered, rather than a combination of correct responses and speed, as with the other categories. The number of days that participants engaged with the program did not differ between groups, although there was considerable variability within groups. Notably, some participants in each of the expectation conditions even continued with the *Activate* program long after they had completed the trial (we did not include data from sessions completed beyond the 5-week trial in our analyses).

Participants who completed the control training (i.e., Sudoku and n-back games) reported being highly engaged with the program, but did not improve on these exercises. Interestingly, participants assigned to the Sudoku and n-back exercises reported that they tended to select a larger number of “hard” Sudoku grids over time, indicating motivation to challenge themselves further. Unfortunately, due to issues with database design, we were unable to analyze performance of the n-back exercises on the basis of level (i.e., 2-vs. 3-vs. 4-back task) here.

We examined amount of logged playtime for both the *Activate* and control games (see Figure 3 and supplemental files) because this was the closest comparison we could make between the two programs. Note, however, that we cannot be certain that the amount of time recorded for the control games consistently reflected engagement in the task, rather than the game remaining open in a web browser after a participant finished, due to reported glitches with the auto-logout. This may explain the relatively large variance for several of the control games. This issue did not appear to exist for the *Activate* games.

Participants assigned to the high expectation priming conditions spent more time on one of the four *Activate* games, compared to their counterparts who received low expectation priming. This suggests that participants in the high expectation conditions may have been more motivated to participate in the program, and may have contributed greater effort throughout the training sessions, at least on some games, compared to those in the low expectation conditions. Nevertheless, lack of significant differences on the training tasks suggests that any possible effects of motivation and effort were minimal, and did not meaningfully contribute to performance.

### (Limited) Transfer Effects Cognitive Function

Although participants provided positive feedback, we did not find any effect of expectation priming or of the CCT program on our transfer measures of cognitive or psychosocial function (see supplemental files). There are several possible reasons.

We recruited a self-selected sample of relatively high-functioning older adults, some of whom who were concerned about their cognitive health and likely to partake in cognitive training on their own. In this regard, our sample is representative of CCT subscribers. Had we recruited more low-functioning older adults or people with psychiatric conditions (e.g., as in Morimoto et al., 2016), we might have found transfer of training. Some promising findings have been reported in people with Mild Cognitive Impairment (MCI), although even in that group the literature is inconsistent (e.g., Butler et al., 2018). One recent high-quality study reported improved memory and strategy use in people with amnesic MCI following eight weekly sessions of group-led cognitive training (Belleville et al., 2018). Nevertheless, there is currently no conclusive evidence that cognitive training can prevent or delay the onset of MCI or other forms of dementia (National Academies of Sciences, 2017).

Although an assessment of strategy use was beyond the scope of this study, it is possible that some participants developed or obtained (e.g., from a friend or family member) more effective strategies on how to complete the games. For example, at post-assessment, a number of participants reported repeating items to be remembered out loud, assigning names to specific repeated stimuli, or placing the cursor in strategic locations in anticipation of a response. This may have limited the benefits of the cognitive training. Similarly, because a large portion of the training was completed from home, we were unable to control for environment during home training sessions, or monitor the potential use of external aids (e.g., other computing software) to complete the Sudoku puzzles or n-back trials.

### Expectation Priming

Regarding the *Activate* training, one possibility for the relative lack of an expectation effect is the subtlety of our priming messages. Although ratings on the EAS appeared to change consistently in the predicted direction in hypothetical and anticipatory scenarios, the messages might be insufficient to alter perceptions when participants have actually undergone an intervention. We elected not to use a more overt or repeated priming strategy to avoid divulging the nature of our expectation manipulation. However, measuring and evaluating the potential influence of expectations on outcomes is difficult, and the most appropriate approach remains debated (Green et al., 2019); it is therefore possible that using other forms of media (e.g., verbal messages or pre-recorded videos) would provide more potent expectation priming. Despite these potential considerations, a recent study using different expectation priming and assessment methods also found little effect of an expectation manipulation (Tsai et al., 2018). One thing to keep in mind in future work is that participants can be quite sophisticated in their ability to recognize or infer the subtleties of experimental design: Many of ours were actively engaged in learning about cognitive aging in their leisure time and participated in research studies as a way to remain informed and involved. Consequently, many stated at the outset or at follow-up that training effects can be nuanced, and that both achieving and measuring transfer can be difficult.

### CCT Design

Although we found no significant expectation or training effects on cognition here, we might have under other conditions. For example, it is possible that longer periods of training would lead to measurable effects. The scientific community has yet to agree on an optimal timeframe and intensity of training (i.e., “dose”; Rabipour, Miller, Taler, Messier, & Davidson, 2017). Nevertheless, the creators of *Activate* agreed that our training schedule and timeline were reasonable, reaching the minimal 1000 minutes of training recommended to commercial subscribers to detect performance changes. Moreover, increasing either of these parameters (i.e., length or frequency of training) would also increase the risk of participant dropout. Offering booster sessions at a pre-determined time interval (e.g., several months or a year following completion of the program) might help maintain or even enhance training effects (Rebok et al., 2014). Nevertheless, this solution is unlikely to have altered outcomes in the present study given the lack of any immediate benefits. Moreover, although we conducted a 6-month follow-up, we were unable to analyze these data due to low response rate. Such low response rates suggest that few participants would have returned for booster sessions.

Our null findings may also have resulted from using sub-optimal transfer measures. Although we selected our transfer tasks carefully, based on evidence from the literature as well as consensus with the creators of the *Activate* program – researchers and medical professionals at Yale University – it is possible that the behavioural measures were not refined enough to detect small behavioural effects or changes at the neurological level. For this reason, we have begun a follow-up trial of the *Activate* program using identical training and control conditions, but with different behavioural tasks as well as electrophysiological outcome measures. In addition, as with all subjective measures, it is possible that responses on our self-reports of expectations, subjective memory, and wellbeing were unrepresentative (e.g., if participants simply told us what they believed we wanted to hear; i.e., “demand characteristics”). However, reported expectation ratings replicated our previous findings, and the subjective questionnaires we administered are commonly used in studies of cognition, aging, and CCT.

Finally, although we conducted an *a priori* power analysis to determine the minimal required sample size, any effects of this training could be smaller than anticipated and therefore would be more easily discernible with a larger sample size. Increasing the number of participants would enable further exploratory analyses, examining the potential influence of individual factors (e.g., differences in baseline performance, mood, subjective perceptions of performance and function, etc.) in mediating training outcomes. Nevertheless, collapsing across expectation conditions permitted us to evaluate program-specific effects with a larger sub-sample; these effects were non significant.

## Conclusions

The present study is the first to directly examine expectations of CCT outcomes, including neuropsychological and psychosocial transfer measures, in the lab setting. By priming participants to have either high or low expectations of training at the outset, we sought to tease apart effects of training from those of expectations of CCT in older adults.

Moreover, we incorporated many of the best practices recommended for high quality intervention research. Crucially, studies rarely account for psychological influences such as expectations of outcomes (Green et al., 2019), a factor we directly evaluated and manipulated experimentally. Examining such factors is particularly important for clinical populations, where placebo effects are known to affect outcomes (Benedetti, Carlino, & Pollo, 2011) and where participants may have more to lose compared to healthy populations. Interestingly, evidence suggests that the placebo effect is weaker, and potentially non-existent, in people with certain forms of dementia (e.g., Alzheimer’s; Benedetti et al., 2006). Expectancy-related placebo effects may therefore represent less of a confound in cognitive training trials in this population.

Our findings replicate previous reports that older adults have high expectations of CCT outcomes (Rabipour & Davidson, 2015). Feedback from our participants further supports the importance of credibility on subjective experience, including engagement with the program. Nevertheless, our 5-week trial of a commercially available CCT program indicated little effect of training program, or of expectations, on cognitive performance or subjective perceptions of cognition and wellbeing. Our findings may reflect the specific parameters of this study, and do not imply that all CCT programs – even other versions of *Activate* – are ineffective. Nevertheless, these results join with others in suggesting caution regarding the effectiveness of such short-term computerized programs on cognitive and behavioural outcomes (e.g., Kable et al., 2017; Wentink et al., 2016; for a review, see Simons et al., 2016).

Given the widespread appeal of CCT as an affordable and accessible form of cognitive enhancement in both healthy and clinical populations, understanding the potential impact of expectations and of program design is imperative to assessing the practical value of such interventions. Thus, although we found few effects of expectations here, future studies of cognitive intervention should account for participant expectations using a validated tool such as the EAS. Optimizing intervention design and harnessing the potential positive effects of expectations can also be useful in the home, clinic, or other settings. Eventually, this understanding may inform the development of more effective approaches towards enhancing cognitive function and optimizing clinical interventions for the treatment or prevention of cognitive impairment.

## Author Contributions

S.R. and P.S.R.D. designed the study with help from M.P. S.R., C.M., and J.C. collected the data. S.R. and P.S.R.D. analyzed the data with help from C.M., J.C., M.O.G.G., and A.P. S.R. and P.S.R.D. wrote the manuscript. All authors reviewed and approved the final version of the manuscript.

## Acknowledgements

We thank Bruce Wexler and Sean O’Leary for granting us free access to and support for the *Activate* program. We also thank Bruce Wexler, Sarah Morimoto, and Morris Bell for providing feedback on our experimental protocol and analysis plan, as well as Veronika Huta for guidance on our statistical analyses. We thank Bruce Wexler for helpful comments on an earlier draft of this manuscript, as well as Charles Collin, Andra Smith, Vanessa Taler, and Louis Behrer for thoughtful feedback on the study design and analysis. We thank Zachary Ball, Lara Geinoz, Victoria Hall, Alix Hill, Katia Giguère Marshal, Awais Rahman, Raphaëlle Robidoux, Zak Sanaye, Jonathan Tran, and Allison Walsh for help with data collection and verification. We also acknowledge the Natural Sciences and Engineering Research Council (NSERC, Discovery Grant), Ontario Graduate Scholarships, and Fonds de Recherche Québec – Santé for their support of this work.

## Supplemental Files

### Supplemental Methods

We included subjective measures in addition to our objective tests to permit a more comprehensive analysis of potential training-related improvements on untrained tasks and general functioning.

#### Psychosocial Indices

#### Multifactorial Memory Questionnaire (MMQ)

The MMQ (Troyer and Rich, 2018) is a meta-memory questionnaire that includes scales of contentment or satisfaction (i.e., feelings regarding memory performance), ability (i.e., self-appraisal of memory capabilities), and strategy (i.e., reported frequency of memory strategy use). Self-report of memory may be an important predictor of cognitive decline (Reid and Maclullich, 2006). Nevertheless, both healthy older adults and those with memory disorders can report inaccurate beliefs about memory and aging – which may reflect negative cultural stereotypes – and poor evaluation of their own memory (Troyer and Rich, 2002). Such inaccuracies may lower expectations, reduce e?ort on everyday tasks involving memory, and lead to impaired memory performance.

#### Pittsburgh Sleep Quality Index (PSQI)

The PSQI is a brief scale measuring sleep quality. Studies suggest that sleep quality may deteriorate in healthy older adults, independent of medication use or the presence of other physical or psychological conditions (Buysse et al., 1991; Nebes et al., 2009). Poor quality of sleep has been shown to associate with greater risk of cognitive decline and dementia (Spira et al., 2014).

#### Subjective Age Questionnaire

We included a measure of subjective age, adapted from (Hughes et al., 2013), which asks participants to rate their subjective age in years on four categories: i) general age; ii) age with respect to wellbeing; iii) physical age; and iv) cognitive age. Studies suggest that older adults feel older after immediately performing memory tests (Hughes et al., 2013), and that perceived age may correlate with cognitive function (Stephan et al., 2016). We sought to determine whether this pattern would persist in our sample or, conversely, whether perceived age would decrease after completing the cognitive intervention. We used a computerized version of the scale, asking participants to indicate their perceived age in each category by dragging the cursor to the appropriate location on a line denoted only by a “0” and “120” on either end. The relative location of the cursor determined subjective age.

#### Patient Health Questionnaire (PHQ-9)

The PHQ-9 is a brief questionnaire that assesses the presence and severity of depression. PHQ-9 reports correlate with mental and general health, as well as social and physical functioning (Kroenke et al., 2001). We included the PHQ-9 in our final 71 participants based on advice from our colleagues, as a more sensitive substitute for the Center for Epidemiological Studies-Depression (CES-D) scale (Lewinsohn et al., 1997).

#### Social Engagement Questionnaire (SEQ)

The SEQ measures indices of social engagement, including social activity, social network size, and social support (Krueger et al., 2009). We included this measure in our final 53 participants based on feedback from our colleagues, and slightly adapted the form under the guidance of Kristin Krueger, who initially developed the scale.

#### Short Form Health Survey (SF-36)

The SF-36 measure of health-related QOL is reliable in older adults (Hayes et al., 1995). The questionnaire categorizes the items into nine scales, including physical functioning, physical health problems, emotional problems, vitality, mental health, social functioning, bodily pain, general health, and health transition.

#### Feedback on Perceived Experience

At the post-training assessment, we asked participants to complete a satisfaction questionnaire for feedback regarding their perceived experience during the intervention. On a scale from 1-7 (1 = “very strongly disagree,” 2 = “strongly disagree,” 3 = “disagree,” 4 = neutral/“neither agree nor disagree,” 5 = “agree,” 6 = “strongly agree,” 7 = “very strongly agree”), participants rated the degree to which they found the experience: i) enjoyable; ii) challenging; iii) frustrating; iv) engaging; v) boring; vi) motivating; and vii) satisfying.

### Supplemental Results

#### Psychosocial Transfer Effects

Repeated measures MANOVA examining subjective reports of psychosocial data across program and expectation condition revealed neither a main effect of time, program, or expectation priming (F_(5,77)_≤2.11, p≥.073, BF10≤.208), nor any interactions between them (F_(5,77)_≤1.81, p≥.122, BF10≤.364; Figure 6).

#### Subjective Experience

The majority of participants (70/87=80%) believed their assigned program was genuinely training their brain. Interestingly, the proportion of participants with this belief was larger, albeit not significantly, in the active control condition (38/43=88%) compared to the Activate intervention (32/44=73%; X^2^ =4.7, p=.066).

Participants provided largely positive feedback and reported that they would recommend the program (Figure 7a), regardless of experimental condition (77/87=89%; *X*^2^ =2.45, ns). MANOVA of participant feedback across expectation priming and CCT program revealed a marginal effect of program (Wilk’s λ=.838, F_(7,77)_=2.12, p=.051, η_p_^2^=.162), but not of expectation priming (F_(7,77)_=.939, p=.482) or an interaction (F_(7,77)_=.714, p=.66), on subjective experience. After correcting for multiple comparisons, participants in the active control group (i.e., training on Sudoku and n-back tasks) reported significantly higher enjoyment (F_(1,83)_=7.48, p=.008, η_p_^2^=.083) and lower boredom (F_(1,83)_=9.42, p=.003, η_p_^2^=.102) of the program.

Perhaps more interestingly, MANOVA of feedback ratings revealed that believing the program genuinely trained the brain significantly influenced subjective experience (Wilk’s λ=.751, F_(7,79)_=3.75, p=.001, η_p_^2^=.25; Figure 7b). Specifically, after correcting for multiple comparisons, people who perceived their assigned program as having genuinely trained their brain rated higher enjoyment (F_(1,85)_=13.47, p≤.0001, η_p_^2^=.137) and engagement (F_(1,85)_=12.42, p=.001, η_p_^2^=.127) with their assigned program, and lower boredom (F_(1,85)_=18.70, p<.0001, η_p_^2^=.180). Note that, due to an unequal distribution of “believers” vs. “non-believers” and, in particular, low number of “non-believers”, we collapsed “belief” (i.e., perception of training) across experimental conditions to perform these analyses.

#### Long Term Outcomes

About half of participants completed at least part of the 6-month follow-up assessment (n=53 responses on the psychosocial questionnaires; n=38 responses on the EAS, final feedback questionnaire, and informal follow-up interview). Due to low power, we elected not to analyze these outcomes.

### Supplemental Discussion

#### Psychosocial Function

We did not detect any changes in self-reported measures of cognitive performance, psychosocial wellbeing, or health-related QOL. This finding may represent the – perhaps un-representative – nature of our participant sample, which may have comprised particularly high functioning older adults. Notably, the older adults we recruited were highly educated, independent, and motivated members of the Ottawa community. Many participants demonstrated considerable insight into their own behaviour as well as our study design and evaluation of the program. MoCA scores suggest that our participants had high global cognition, with means in all experimental conditions above the clinical cutoff of “26/30”. Moreover, participants tended to have high subjective perceptions of cognition, wellbeing, and QOL at baseline. Variability in these measures, however, may moderate the effects of cognitive training (e.g., people with low ratings of affect or QOL may benefit from CCT to a greater extent; Morimoto et al., 2016). Although we did not find large changes in these outcomes, future studies with greater sample sizes may consider analyzing these variables as mediators or predictors rather than direct outcomes, as we did here.

#### Subjective Perceptions

Participants enjoyed both the *Activate* and control (Sudoku/n-back) program. Interestingly, participants seemed to enjoy the Sudoku and n-back exercises more than the *Activate* games, and more participants assigned to those exercises believed the program was genuinely training the brain compared to participants who completed *Activate* training. Moreover, we found greater enjoyment and engagement, as well as lower boredom in participants who believed the program had genuinely trained their brain. This finding suggests that, similar to other contexts (e.g., in medicine; Gollub and Kong, 2011; Wampold and Imel, 2007), belief in the capacity of a CCT program to achieve its intended goal (i.e., train the brain) can significantly impact behavioural and other outcomes. Although we did not have a large enough sample of “believers” vs. “non-believers” across our experimental conditions to probe this effect further, future studies may wish to examine how such retrospective perceptions of a program’s effectiveness may interact with other outcomes.

